# Experimental Genetics Validation of *Plasmodium falciparum* Gametogenesis Essential Protein 1 (GEP1) as a Transmission Blocking Target

**DOI:** 10.1101/2025.05.28.656680

**Authors:** Frederik Huppertz, Milagros Siebeck Caturelli, Lina Lehmann, Florian Kurth, Alexander G. Maier, Kai Matuschewski

**Affiliations:** Dept. of Molecular Parasitology, Institute of Biology, Humboldt University, 10115 Berlin, Germany; Research School of Biology, The Australian National University, Canberra 2601, Australia; Department of Infectious Diseases and Critical Care Medicine, Charité - Universitätsmedizin Berlin, Berlin, Germany; Centre de Recherches Médicales de Lambaréné, 242 Lambaréné, Gabon

**Keywords:** malaria, *Plasmodium*, transmission, transport protein, gametogenesis, exflagellation

## Abstract

Transmission of *Plasmodium* parasites to the *Anopheles* vector critically depends on swift activation of mature gametocytes upon entry into the mosquito midgut. Induction of gametogenesis requires two simultaneous stimuli, a temperature drop and xanthurenic acid. Previous work in the murine malaria model *Plasmodium yoelii* identified a protein, termed gametogenesis essential protein (*GEP1*), with a suggested role in xanthurenic acid-dependent activation of gametes. Here, we present an experimental genetics characterization of *GEP1* in the human pathogen *Plasmodium falciparum*. Using CRISPR-Cas9 gene editing we generated *PfGEP1* loss-of-function lines and analyzed their progression until gametocyte maturation. We show a complete defect in both male and female gametogenesis caused by disruption of *PfGEP1*. *Pfgep1(-)* gametocytes do not produce gametes when activated with xanthurenic acid or a drop in temperature. This defect could not be overcome by the phosphodiesterase inhibitor Zaprinast, which induces gametogenesis. We also explored *GEP1* haplotypes in *P. falciparum* parasites circulating in endemic regions and show the presence of two non-synonymous SNPs, resulting in V241L and S263P mutations, in 12% and 20% of 49 sentinel samples, respectively. Together, our data indicate that GEP1 plays a central role in the gamete activation process independent of xanthurenic acid and validates *Pf*GEP1 as a promising transmission blocking target.

## Introduction

Malaria remains the most important arthropod-borne infectious disease with an estimated 263 million infections and 597,000 deaths per year (WHO 2024). *Plasmodium* parasites, the causative agent of malaria, follow a complex developmental program in the vertebrate host, where they can cause life-threatening disease, and the *Anopheles* vector, where sexual recombination takes place. Accordingly, maturation of the sexual precursor cells in the human blood, termed gametocytes, represents a potential point of attack for transmission intervention strategies, which are considered pivotal for malaria elimination (Birkholtz et al. 2022)

Blood-stage parasites repeatedly infect and replicate inside erythrocytes and eventually commit to sexual stages (Josling et al., 2016). Maturation of *P. falciparum* gametocytes typically occurs over the course of 10-12 days and is commonly divided into five morphologically distinct stages (I-V) (Talman et al., 2004; Baker, 2010). Whereas the early stage I gametocytes are almost indistinguishable from their asexual counterparts, starting from stage II gametocytes increase in volume and begin accumulating hemozoin pigment in a sex-dependent manner. Female gametocytes exhibit condensed hemozoin crystals while male parasites distribute the crystals more scattered throughout the cell. These differences become more apparent as gametocytes develop further. During stages III to V, parasites elongate and give the host cell the characteristic sickle-cell, so-called falciform, shape that the species derives its name from. Ultimately, gametocytes serve the purpose of transmission from the human host back into the mosquito vector.

Once gametocytes are taken up by a mosquito during its blood meal, they are activated by the drop in temperature and a rise in pH, as well as the presence of xanthurenic acid (XA) in the mosquito midgut (Billker et al. 1998; Brochet et al. 2021). How exactly these stimuli translate into intracellular activity is not yet fully understood, but shortly after transmission an intracellular signalling cascade commences, beginning with an increase in cGMP concentration mediated by GCα (Muhia et al. 2001; Kawamoto et al. 1990). cGMP is subsequently used by a protein kinase (PKG) to phosphorylate a multipass membrane protein, termed important for calcium mobilization-1 (ICM1), which was suggested as a Ca^2+^ channel that mediates Ca^2+^ release from internal storages (Balestra et al. 2021). Phosphoinositide-specific phospholipase C (PI-PLC) is known to be required for this Ca^2+^ mobilization (Raabe et al. 2011), but the interactions between PI-PLC, ICM1 and Ca^2+^ remain unresolved. Downstream of the Ca^2+^ release, several calcium-dependent protein kinases (CDPKs) and calcineurin regulate egress events. There is some overlap between male and female gametogenesis, including the reliance on CDPK1 to mediate egress from the host cell (Bansal et al. 2018). In contrast, CDPK2 appears to function only in male gametocyte egress (Bansal et al. 2017). Finally, eight microgametes are released by the male gametocyte that can fertilize a macrogamete formed by the egressed female gametocyte.

Recently, a candidate transport protein, termed gametogenesis essential protein 1 (GEP1), was identified in the rodent malaria model parasite *Plasmodium yoelii* as an essential component of gametocyte activation (Jiang et al. 2020). GEP1 colo^-^ calizes with GCα and was suggested to be needed for its activity. Parasites deficient in *PyGEP1* did not exhibit increased cGMP concentrations, which is typically detected upon stimulation with xanthurenic acid, indicating that GEP1 may function upstream of cGMP in the signalling cascade (Jiang et al. 2020). This defect could not be bypassed with the phosphodiesterase (PDE) inhibitor Zaprinast, which has been shown to trigger gametogenesis in wild type parasites (McRobert et al. 2008). Upon Zaprinast inhibition the basal activity of GCα eventually increases cGMP levels above the threshold without further activation, since the counteracting PDE activity is inhibited. The lack of Zaprinast-induced gametogenesis in *PyGEP1*-deficient parasites supports the notion that GEP1 is needed for this basal activity of GCα. While the exact role of GEP1 in gametogenesis remains unknown, the complete defect of *gep1(-)* parasites in gamete egress qualifies GEP1 as an attractive target to prevent parasite maturation in the mosquito vector. Whether *GEP1* defects reproduce in *P. falciparum* and, hence, qualifies as a candidate drug target awaits experimental genetics confirmation.

*PfGEP1* encodes a protein of 960 amino acids with 17 predicted transmembrane domains and appr. 250 amino acids reaching into a non-cytoplasmic compartment at the N-terminus (Fig. 1A). Its classification as a transporter was inferred by weak similarity to a family of Na^+^-neurotransmitter symporters that transport a wide array of substrates, ranging from neurotransmitters to amino acids (Amara and Arriza 1993). Accordingly, it was originally labelled NSS2 (Martin 2020). Sequence alignment of *Pf*GEP1 with its orthologs in the *Plasmodium* genus revealed conservation within the *Laverania* subgenus and 80-90% similarity to genes found in the *Plasmodium*, *Vinckeia*, *Hepatocystis* and *Haemamoeba* subgenera (Fig. 1B). In *Hepatocystis*, two orthologs were identified on two separate chromosomes corresponding to half the *Pf*GEP1 protein each. These most likely represent sequence mis-assembly in the draft *Hepatocystis* genome (Aunin et al. 2020). Outside of the genus *Plasmodium*, no orthologs of GEP1 can be identified based on sequence similarity.

**Figure 1:**
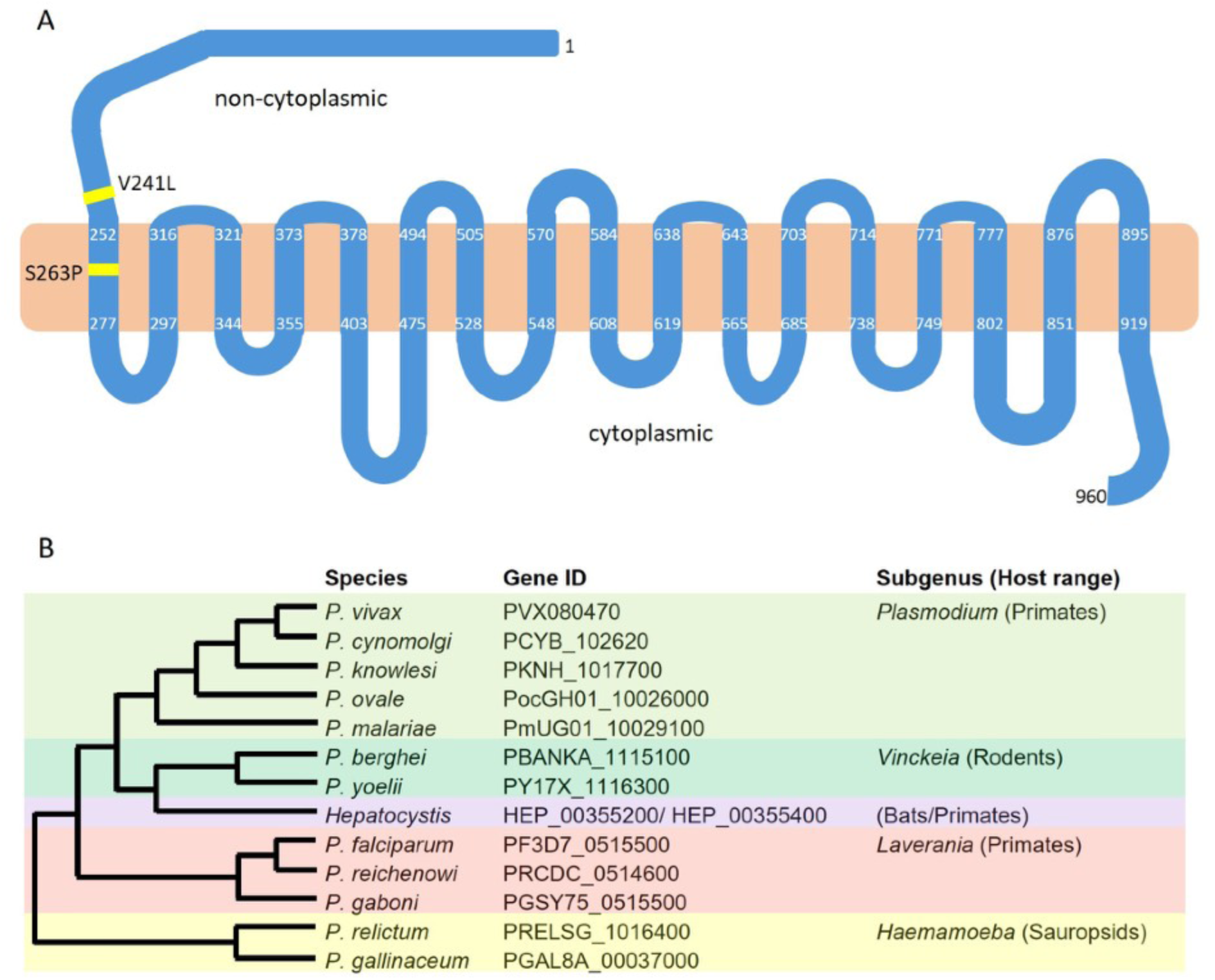
Structural prediction and orthology of *PfGEP1*. **A** *Pf*GEP1 encodes for 960 amino acids, including 17 predicted transmembrane (TM) spans. The residues defining the TMs are indicated. The amino-terminus of appr. 250 amino acid residues and the shorter carboxyterminus are predicted to be noncytoplasmic and cytoplasmic, respectively. Two single nucleotide polymorphisms (SNPs), V241L and S263P, have been identified (yellow lines). **B** Representation of *GEP1* orthologs across the genus *Plasmodium*. Shown are *Plasmodium* species, gene identifier, subgenus and host taxa.

Two non-synonymous single nucleotide polymorphisms (SNPs) in the *PfGEP1* gene were reported thus far (Mu et al. 2003; Bustamante et al. 2012). The V241L polymorphism was identified in a study searching for markers of chloroquine/quinine resistance, but no association was observed for this allele and altered drug efficacy (Mu et al. 2003). The V241L SNP, as well as a second SNP, S263P, was later reported to be associated with reduced *in vitro* susceptibility to artemether in samples collected in Nigeria (Bustamante et al. 2012), but a functional link between these SNPs and drug resistance remains to be determined.

Here, we report a complete lack of gametogenesis in parasites deficient for *PfGEP1* in the human pathogen *P. falciparum*. We found this defect to appear independently of XA and to not be reversible by addition of the phosphodiesterase inhibitor Zaprinast, lending further support for the notion that GEP1 is necessary for basal activity of GCα rather than being a receptor for the external stimulus. This observed arrest in gamete activation in a human malaria parasite qualifies *Pf*GEP1 as a candidate transmission blocking target. We also assessed the prevalence of the known SNPs in the *PfGEP1* gene in a sample collection obtained from travelers returning from African countries with malaria.

## Results

### Expression profiling of *PfGEP1* in cultured erythrocytes

We initiated our study by assessing the steady-state transcript profiles of *PfGEP1* by RT-qPCR of selected asexual and sexual blood stages relative to three housekeeping genes, namely fructose-bisphosphate aldolase (PF3D7_1444800), seryl tRNA synthetase (PF3D7_0717700), and HSP70 (PF3D7_0818900). This analysis revealed low expression during asexual blood stages and a marked increase throughout all gametocyte stages (Fig. 2A). Interestingly, the highest expression was found in early gametocytes (day 4), and expression decreased towards mature gametocytes. Together, the expression data indicate that *PfGEP1* likely plays a minor role during asexual blood infection *in vitro* and has an important function during gametogenesis, but not necessarily during the final stages of gamete activation.

**Figure 2.**
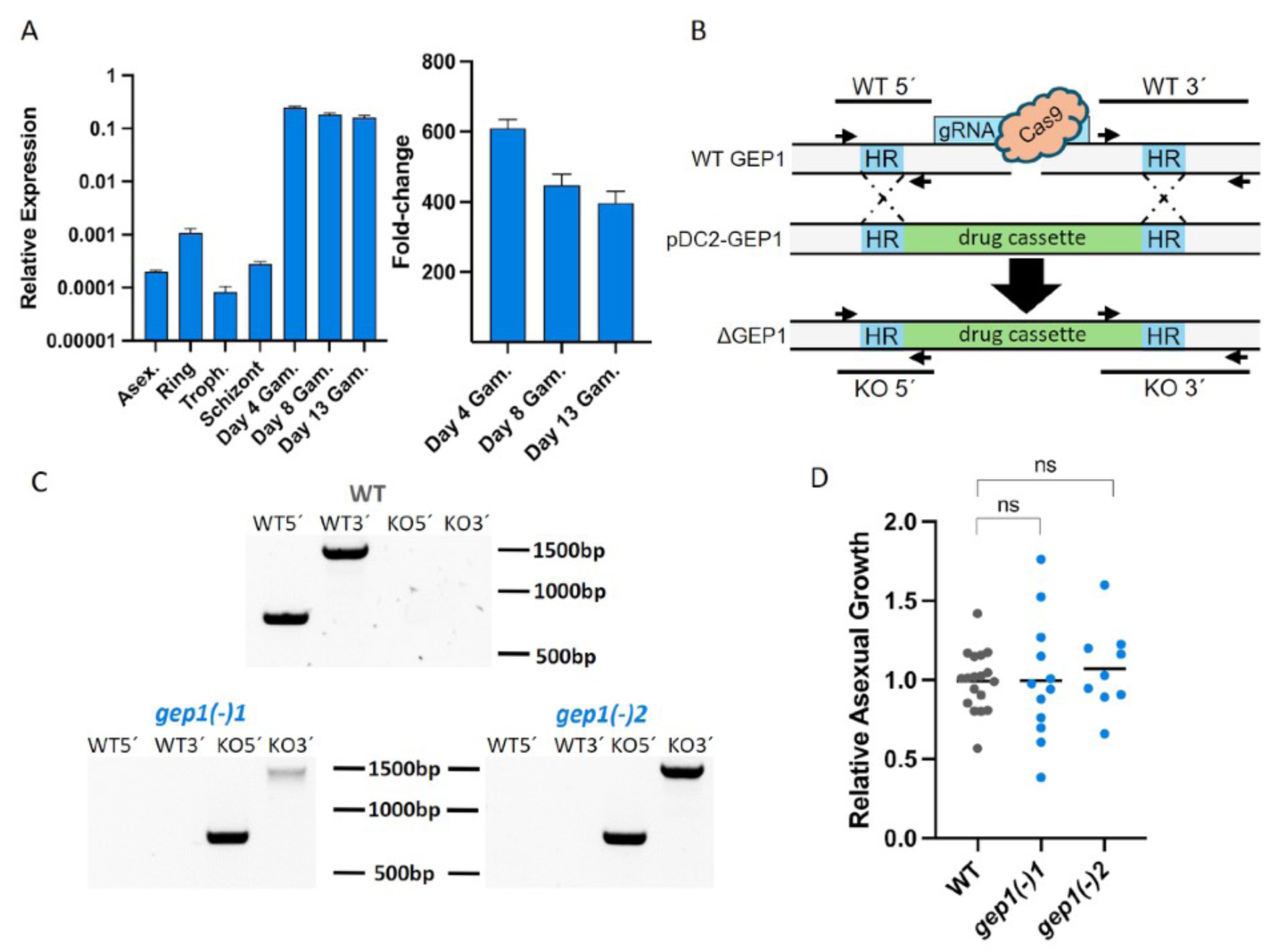
Expression profiling and targeted deletion of *PfGEP1*. **A** RT-qPCR analysis of *PfGEP1* expression during asexual and sexual blood infection. Shown are relative expression levels (ΔΔCt values) compared to three house^-^ keeping genes (seryl tRNA-synthetase, fructose bisphosphate aldolase and *HSP70*) (left). Mixed asexual stages, synchronized ring stages, synchronized trophozoites, synchronized schizonts, and gametocytes were harvested 4, 8, and 13 days after induction and the fold-change of *PfGEP1* expression in gametocytes in comparison to mixed asexual stages determined (right). Data are from one sample set done in three technical replicates. **B** Disruption strategy to generate *gep1(-)* parasite lines using a Crispr-Cas9-based approach. Shown are the wild-type (WT) genomic locus, the targeting plasmid (pDC2-GEP1) and the predicted recombinant locus after homologous recombination (ΔGEP1). Homology regions (HR, blue), the drug resistance cassette (green) for positive selection, diagnostic primers (arrows) and PCR products (lines) are indicated. **C** Diagnostic PCR to verify successful *PfGEP1* disruption in two separate cell lines. Primer combinations and PCR products as indicated in B. **D** Asexual growth of *gep1(-)* cell lines relative to NF54 wild type parasites. Growth was monitored over 48 hours and shown normalized to NF54 growth rates. n.s., non-significant (*p*>0.05; t-test, three biological replicates with three technical replicates each).

### *PfGEP1* is dispensable in asexual blood stages

Next, two CISPR/Cas9-based plasmids were generated to disrupt the *PfGEP1* gene in cultured *P. falciparum* parasites (Fig. 2B). Parasites were visible 26 and 49 days after transfection with the respective plasmids. Successful disruption of *PfGEP1* was confirmed by diagnostic PCR. The parasite populations originated from two independent guide RNAs ini the CISPR/Cas9 targeting plasmids and appeared isogenic. Accordingly, we selected two respective clonal lines, termed *gep1(-)-1* and *gep1(-)-2*, which exhibited the desired genotype, validating successful *PfGEP1* deletion (Fig. 2C).

Clonal parasite lines were examined for asexual blood replication (Fig. 2D). In these growth assays no differences of *gep1(-)-1* or *gep1(-)-2* growth were observed compared to NF54 wild type (WT) parasites.

### *PfGEP1* does neither affect gametocyte commitment nor maturation

To characterize sexual differentiation of *gep1(-)* parasites, gametocyte commitment was induced and ring stage parasites (sexually and asexually committed) were counted. Four days later, when asexual parasites had been removed, early gametocyte parasitemia was assessed and the commitment rate calculated (Fig. 3A,B). While commitment differed between repeats, no significant difference in conversion rate between WT and *gep1(-)* parasites was observed (Fig. 3B). We next assessed the rate of day 4 gametocytes that reached maturity on day 10 (Fig. 3A,C). The majority of gametocytes matured, and there was no difference between WT and *gep1(-)* parasites. We also enumerated the ratio of male and female gametocytes (Fig. 3D). Again, there was no apparent difference between WT and *gep1(-)* parasites. Together, we show that absence of *Pf*GEP1 does not interfere with sexual differentiation of cultured *P. falciparum* parasites.

**Figure 3.**
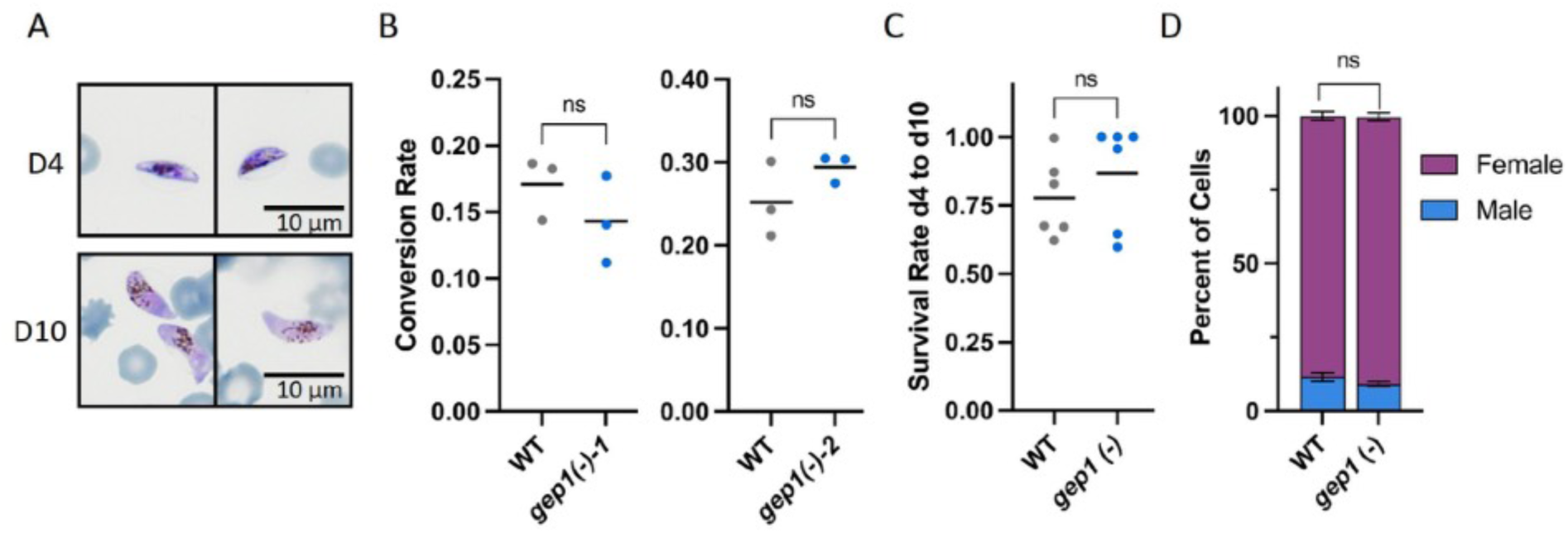
Disruption of *PfGEP1* does not affect gametocyte development. **A** Developing (top) and mature (bottom) gametocytes produced by *gep1(-)* parasite lines. **B** Comparison of gametocyte commitment between *gep1(-)* and WT parasites. WT and *gep1(-)*-1 and *gep1(-)*-2 parasites, respectively, displayed similar gametocyte conversion rates. Parasitemias quantified in >2,000 erythrocytes, each graph represents one biological replicate with three technical replicates. **C** Survival rate of day 4 gametocytes to reach full maturity. *Gep1(-)* and WT parasites displayed similar rates of full gametocyte maturation. Parasitemias quantified in >2,000 erythrocytes, shown are two biological replicates with three technical replicates each. Rates above 1 were set to 100% survival. **D** Sex ratio of female and male gametocytes. No differences in the proportion of male-to-female ratios were detected between mature *gep1(-)* and WT gametocytes. >100 parasites assessed per culture, three biological replicates. n.s., *p*>0.05 (t-test).

### *PfGEP1* is essential for male gametogenesis

To mimic host switch during the mosquito blood meal, we induced exflagellation in mature gametocytes with or without addition of 100 µM xanthurenic acid (XA) (Fig.4). NF54 wildtype parasites produced exflagellation centres under both conditions. Without XA, WT parasites displayed substantial residual activation, which could be increased 10-fold by XA addition (Fig. 4A,B). Strikingly, *gep1*(-) parasites displayed a complete absence of exflagellation (Fig. 4A), and this defect was independent of XA addition (Fig. 4B).

**Figure 4.**
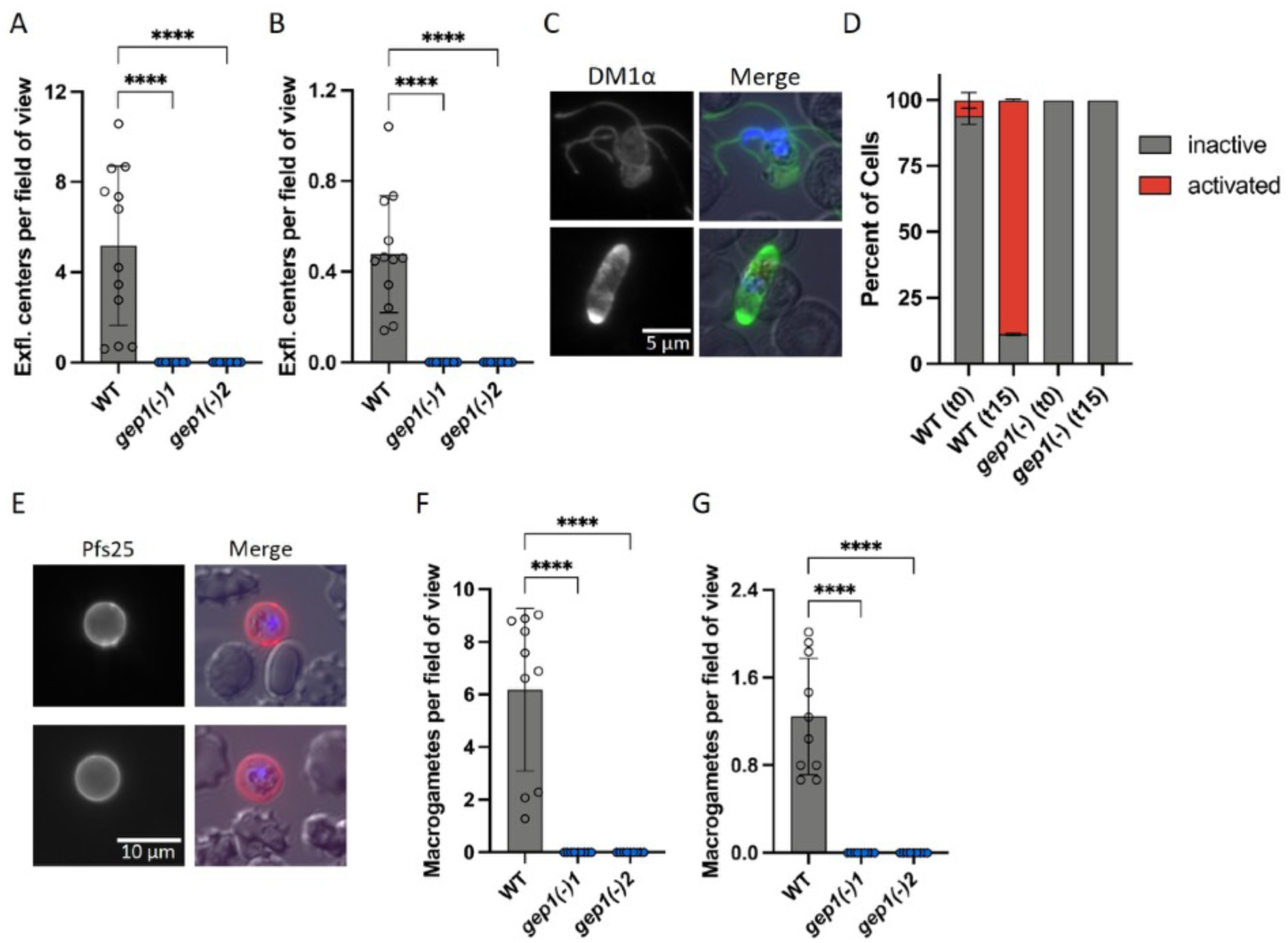
*gep1(-)* parasites display a complete arrest prior to becoming male or female gametes. **A,B** Enumeration of exflagellation centres per field of view (400x magnification) in the presence (A) or absence (B) of xanthurenic acid. 25 fields of view assessed per replicate, shown are four biological replicates with three technical replicates each. **C,D** Mature gametocytes were fixed before or after 15 minutes with XA activation, stained with a DM1α antibody (green) and DAPI (blue), and the ratio of activated (red) and non-activated gametocytes (grey) were quantified (right) (>100 parasites quantified, three technical replicates). Shown are representative images of an activated (top) and a non-activated (bottom) wild type gametocyte (left). **E** Mature gametocytes were activated for two hours and macrogametes stained with a *Pf*s25 antibody (red) and Hoechst (blue). Shown are two representative images of wild type macrogametes. **F,G** Enumeration of macrogametes per field of view (400x magnification) in the presence (E) or absence (F) of xanthurenic acid. 25 fields of view assessed per replicate, shown are three biological replicates with three technical replicates each and an additional biological replicate with only one technical replicate. ****, *p*<0.001 (t-test)

The absence of exflagellation events in *gep1(-)* parasites was confirmed through immunofluorescence assays using a tubulin antibody (DM1α) on cells fixed either immediately or after incubation with XA for 15 min (Fig. 4C,D). As expected, exflagellation centres in WT parasites were readily detected, and quantification revealed appr. 90% activation in WT parasites (Fig. 4D). In non-activated samples, a background activity of appr. 5% was quantified in WT parasites. Notably, the gametocytes visible in the *gep1(-)* samples were neither rounded up nor did they show any other signs of activation. In conclusion, male gamete exflagellation, including XA-independent activation, was completely abolished in the absence of *PfGEP1*.

### *PfGEP1* is essential for female gametogenesis

Activation of female gametocytes was assessed during a prolonged 2-hour activation either with or without XA. To this end, cells were stained with an antibody against the zygote surface marker *Pf*s25 and DAPI (Fig. 4E). Quantification using fluorescence microscopy revealed an average of 6.2 activated female WT gametes per field of view in the presence of XA (Fig. 4F). In the absence of XA, activation of female WT gametes was reduced to 1.2 per field of view (Fig. 4G). In marked contrast and in good agreement with the complete defect in male exflagellation, activated female gametes were completely absent in the *gep1*(-) samples (Fig. 4F,G).

### The *gep1(-)* defects cannot be rescued by the phosphodiesterase inhibitor Zaprinast

We next tested whether the gametogenesis defect of *gep1*(-) parasites can be, at least partially, overcome by addition of a cyclic guanosine monophosphate (cGMP)-specific phosphodiesterase (PDE) inhibitor. Accordingly, mature gametocytes were activated in the presence of 400μM Zaprinast, and assays were again performed with or without addition of 100μM XA (Fig. 5). In good agreement with an increase in cGMP steady state levels, Zaprinast did not modify the number of exflagellation events of WT parasites in the presence of XA, but elevated exflagellation to appr. 4.2 per field of view without XA (Fig. 5A). A similar effect was observed when assessing macrogametes, where the addition of Zaprinast increased the number of gametes produced to 4.5 gametes per field of view without XA (Fig. 5B). In marked contrast, *gep1(-)* parasites remained unable to produce either male or female gametes under any of the tested conditions indicating that cGMP levels in these cells cannot be elevated by a cGMP-specific PDE inhibitor and bypass the critical role of *Pf*GEP1 in gamete activation.

**Figure 5.**
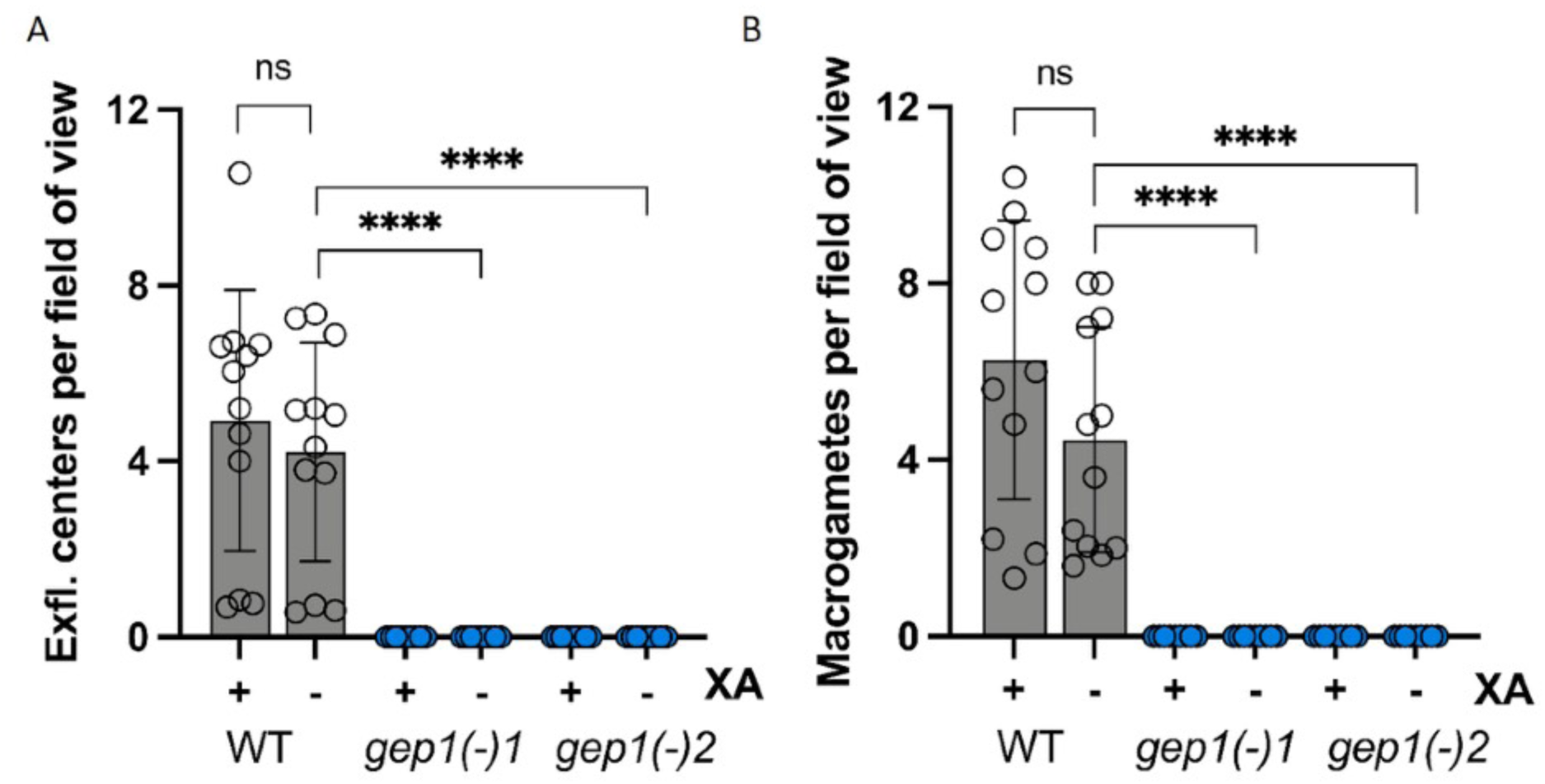
The phosphodiesterase inhibitor Zaprinast does not rescue defects of gamete activation in *gep1(-)* parasites. **A** Quantification of exflagellation centres in the presence or absence of 400 µM Zaprinast. **B** Quantification of macrogametes in the presence or absence of Zaprinast. A/B: 25 fields of view assessed per replicate, shown are four biological replicates with three technical replicates each. ****, *p*<0.001 (t-test) XA-xanthurenic acid

### Genetic diversity of *PfGEP1* in field samples

We finally employed an amplicon sequencing protocol and assessed the prevalence of the V241L and S263P polymorphisms in a collection of 52 clinical samples opportunistically collected from travelers returning from 15 different, mostly Western African, malaria-endemic countries. GEP1 amplicons were successfully generated with a GEP1SNP primer pair for 49 samples (Table 1). Sanger sequencing revealed the V241L allele in 6 samples (12%), while the S263P allele was identified in 10 samples (20%). Notably, three samples (6%) exhibited both SNPs. In our limited dataset, we observed the two SNPs across Sub-Saharan Africa, including Kenya and Rwanda as well Côte d’Ivoire and Gambia. We suggest that *GEP1* SNPs are widespread and moderately frequent.

**Table 1:** Prevalence of *GEP1* SNPs in samples collected from returning travelers.

| Country of origin | Samples | V241L<sup>+</sup> | S263P<sup>+</sup> | V241L<sup>+</sup>/S263P<sup>+</sup> |
| --- | --- | --- | --- | --- |
| Nigeria | 14 | 0 | 2 | 1 |
| Cameroon | 10 | 1 | 1 | 0 |
| Ghana | 4 | 0 | 1 | 0 |
| Togo | 4 | 1 | 0 | 0 |
| Benin | 3 | 0 | 0 | 0 |
| Burkina Faso | 2 | 0 | 0 | 1 |
| CAR | 2 | 0 | 0 | 0 |
| Guinea | 2 | 0 | 1 | 0 |
| Côte d'Ivoire | 2 | 0 | 0 | 1 |
| Sierra Leone | 1 | 0 | 0 | 0 |
| South Africa | 1 | 0 | 0 | 0 |
| Gambia | 1 | 0 | 1 | 0 |
| Rwanda | 1 | 0 | 1 | 0 |
| Congo | 1 | 0 | 0 | 0 |
| Kenya | 1 | 1 | 0 | 0 |
| Total | 49 | 3 | 7 | 3 |

## Discussion

Blocking transmission of *Plasmodium* from the human host to the *Anopheles* vector is one of the main goals of malaria control and remains a research priority. Gametocytes spend most of their maturation time in the bone marrow and frequently escape detection (Joice et al. 2014). The protracted maturation of *P. falciparum* gametocytes over the course of ten days offers a broad window of opportunity for drug interference. Here, we were able to show that loss of function of *PfGEP1* completely blocks gametogenesis of both male and female gametocytes, fully supporting data from the *P. yoelii* murine malaria model (Jiang et al., 2020). Based on the colocalization with GCα, *Py*GEP1 was suggested to interact with GCα and act as a regulator of GCα activity. Our data are in line with the notion that *Pf*GEP1 is necessary for the basal activity of GCα, since the *gep1(-)* defect could not be bypassed with Zaprinast. The basal activity of GCα should be sufficient to elevate cGMP levels above the threshold when PDEs are inhibited, even in the absence of external cues leading to GCα activation (McRobert et al. 2008).

Recently, two additional proteins have been implicated in GCα activity, termed signalling linking factor (SLF) and unique GC organizer (UGO) (Kuehnel et al. 2023). Similar to *gep1(-)* lines, parasites deficient for *SLF* or *UGO* showed impaired gametogenesis. Interestingly, this defect could be bypassed by Zaprinast in *UGO*-deficient parasites, but not in those lacking *SLF* (Kuehnel et al. 2023). Together, it appears that GEP1 and SLF are required for efficient GCα activity in mature gametocytes, either by direct binding or upstream of GCα activation. In contrast, UGO appears to elevate GCα activity and trigger gametogenesis in response to external stimuli in the mosquito midgut, making UGO currently the best candidate for the long-sought XA receptor.

Experimental genetics in *P. yoelii* initially assigned this role to *GEP1* (Jiang et al., 2020). This hypothesis was further supported by structural analysis done on the interactions between GEP1, XA and GCα, which postulated a candidate XA-binding pocket in the GEP1 protein (Zhu et al. 2023). In WT parasites, we observed activation of both male and female gametocytes independently of XA, most likely caused by the drop in temperature and an elevated pH upon removal from cultures resulting in a reduced PDE activity (Kuehnel et al. 2023). Strikingly, this basal level of XA-in-dependent activation was also entirely absent in *gep1(-)* parasites, indicating that direct XA binding, if any, is unlikely the exclusive function of GEP1.

In this study, we analyzed the prevalence of two GEP1 non-synonymous SNPs, V241L and S263P in parasite isolates from patients with malaria. The SNPs were originally described from Nigeria (Bustamante et al. 2012), where the prevalence reached 25%. We found both SNPs in samples from various African countries, and the SNPs were moderately abundant, including several samples that were V241L^+^/S263P^+^. One study suggested that both SNPs might contribute to reduced artemether susceptibility (Bustamante et al. 2012). Given the distinct defect of *gep1(-)* parasites in the final step of *P. falciparum* gamete maturation combined with the remarkably low expression level observed in asexual parasites (López-Barragán et al. 2011, our data), we consider it highly unlikely that *GEP1* mutations confer drug resistance during asexual development. We instead propose that allele diversity in *GEP1* might influence onward transmission to the *Anopheles* vector and future malaria episodes.

In conclusion, our experimental genetics analysis in *P. falciparum* qualifies *Pf*GEP1 as a transmission blocking target. To further examine this potential for targeted drug development, biochemical studies are needed to assign a catalytic and/ or regulatory activity to *Pf*GEP1. Since *Pf*GEP1 is a membrane-spanning protein, the precise localization and cellular compartment will be critical determinants for a better molecular understanding of GPE1 functions. Its annotation as a transporter (Martin, 2020) warrants strategies to identify the GEP1 cargo. Whether GEP1 can be targeted by a drug also depends on the hitherto unknown functional role(s) of the amino-terminus, which contains two closely adjacent non-synonymous SNPs, widespread in endemic parasite populations across Sub-Saharan Africa.

## Methods

### Ethical approval

Blood samples were collected from returning travelers with *P. falciparum* malaria treated at Charité Universitätsmedizin Berlin between 2015 and 2022 within the framework of the Study on Determinants of Malaria Semi-Immunity and Tolerance (DEMIT). The study was approved by the ethics committee of Charité Universitätsmedizin Berlin (EA4/092/21). For parasite cultures O^+^ erythrocytes were purchased as concentrate from the German Red Cross, and human serum (blood groups A^+^/B^+^/ AB^+^) was purchased from Haema AG.

## Parasite maintenance

*P. falciparum* parasites were maintained in RPMI1640 media (PAN Biotech, containing 25 mM HEPES and 2.0g/L NaHCO_3_) supplemented with 480 µM hypoxanthine and 20 µg/ml Gentamicin. Parasites were cultured in medium containing 10% w/v heat inactivated human serum (blood groups A^+^/B^+^/AB^+^, obtained from Haema AG) and in O^+^ erythrocytes (German Red Cross) at a 4% hematocrit at 37°C and slight agitation to prevent settling of the cells. Cultures were gassed with a premade gas mixture of 5.0% CO_2_ and 3.0% O_2_ in N_2_ (Westfalen AG). Parasite growth was monitored by microscopic examination of Giemsa-stained blood smears. Parasites were regularly treated with Sorbitol to maintain synchronicity (Lambros and Vanderberg, 1979).

### RNA extraction and qRT-PCR

Parasites were freed from surrounding erythrocytes through saponin lysis (Christophers and Fulton, 1939). RNA was extracted according to the manufacturer’s protocol (Macherey Nagel). RNA concentration was measured on a Nanodrop ND 1000 (peQLab, Germany), and RNA was immediately used for cDNA synthesis using the Superscript IV Reverse Transcription kit (ThermoFisher) with or without (control) addition of reverse transcriptase. cDNA samples were mixed with Power Sybr Green Master mix (ThermoFisher) according to the manufactureŕs protocol; appr. 2ng cDNA was added for each qPCR reaction. Primers were added at a final concentration of 0,5 µM. The qPCR reaction was performed using a Quantstudio 1 (Thermo-Fisher) with the protocol set as follows: 95°C for 2 minutes, then 40 cycles of 95° for 15 seconds, 56°C for 30 seconds and 60°C for 30 seconds. This procedure was followed by a preset melting curve (60°C for 1 minute ramping up to 95°C at 0.15°C/ second). The resulting Ct values were used to calculate relative expression based on the ΔΔCt method (Livak and Schmittgen 2001) against seryl tRNA-synthetase (*hk1*), Fructose bisphosphate aldolase (*hk2*), and *HSP70* (*hk3*).

### Generation of CRISPR-Cas9 gene disruption plasmids

gRNAs were designed using an online tool (Benchling.com) with the following settings: single guide, 20 bp long, map against 3D7 genome, PAM: NGG (SpCas9, 3 ’side). Two gRNAs with an on-target score >50 and an off-target score >90 were selected (Supplemental Table S1). Primers were designed to amplify two homology regions (HR1/2) of 300-500 bp length around the two gRNA binding sites (Supplemental Table S2). Homology regions were amplified from 20ng NF54 genomic DNA per reaction using DreamTaq Polymerase (Thermo Fisher). Homology regions were inserted into the pDC2-hdhfr-Cas9 plasmid backbone (Lim et al. 2016) through Gibson Assembly (Gibson et al. 2009) to surround the *hDHFR* cassette at the *PspOM*I, *EcoR*I and *Aat*II restriction sites. Successful integration of homology regions was confirmed through analytical digests using diagnostic combinations of restriction enzymes. gRNA oligonucleotides were annealed and inserted at the *Bbs*I restriction site into the pDC2-Cas9-hdhfr backbone containing the homology regions. Successful insertion of the gRNAs was confirmed using Sanger Sequencing (LGC Genomics).

### Transfections

Parasites were transfected with 100μg plasmid resuspended in 15μl TE-buffer (Wu et al. 1995). At least 8 hours after transfection, but before parasites completed their current replication cycle, the medium was removed and replaced with medium containing 4nM WR99210 (Jacobus Pharmaceutical). Medium was changed daily for 10-14 days, then three times a week, and fresh erythrocytes were added weekly. Once parasites emerged, they were analyzed by diagnostic PCR. Recombinant parasite populations with the desired gene deletion were purified by clonal dilution and used for assays.

### Growth assay

Asexual cultures were synchronized twice with sorbitol in a six-hour interval (Lambros and Vanderberg 1979). The next day, the parasitemia of late-stage parasites was calculated by microscopic examination of Giemsa-stained blood films. From each culture, three technical replicates were set up at 0.1% parasitemia. Two days later, parasitemia was assessed by counting >2,000 erythrocytes per culture. Parasitemia of each culture was normalized against the mean parasitemia of the WT reference line NF54 for each replicate.

### Gametocyte culture

Synchronized cultures (Lambros and Vanderberg 1979) were set up at 2% late stage parasitemia at 3% hematocrit (day -3). The day after, half the medium was replaced by new medium (day -2). Parasites were allowed to reach the mature schizont stage and were split to 2% just before merozoite egress (day -1). When committed ring stages were present at the onset of the next cycle, the medium was changed to medium containing 50 mM GlcNAc (day 0). On day 1 of gametocyte development, cultures were treated with sorbitol. Medium was changed daily with GlcNAc medium being used for 6-8 days until no visible asexual parasites remained in the culture. Onward, GlcNAc-free medium was used and media changed daily for at least three more days before gamete egress assays were performed.

### Exflagellation assay

Mature (day 10 onward) gametocytes were briefly centrifuged (800g, 1 minute), and 4µl of infected erythrocytes were resuspended in fresh medium containing either 100 µM xanthurenic acid or a 1:1000 dilution of DMSO as control. This was done while maintaining the parasites at 37°C. Parasites were then incubated for 12 minutes at room temperature before exflagellation centres were counted microscopically. Exflagellation centres were quantified over 25 fields of view (400x magnification) of equally distributed cells.

### Macrogamete assays

Activation of female gametocytes was induced similar to the male cells, but incubated for 2 hours at room temperature, after which cells were briefly centrifuged (800g for 1 min.) and stained with mouse anti-*Pf*s25 (1:500 in PBS) and Hoechst (1:1,000 in PBS) for 30 min. at 4°C. Cells were washed once with PBS and then imaged using a Zeiss AxioImager fluorescence microscope. Rounded, DAPI- and Pfs25-positive cells were counted as macrogametes in 25 fields of view (400x magnification) of equally distributed cells.

### Staining of activated gametocytes

Activated mature gametocytes were carefully layered on a coverslip, air-dried and fixed in MeOH at -80°C for 10 minutes. Cells were permeabilized with 0,05% Saponin in 1% BSA/PBS for 30 minutes at room temperature and washed three times with 0,01% Saponin in 1% BSA/PBS (blocking solution) before primary antibody (anti-tubulin, mouse DM1α, Sigma Aldrich T6199) was added (1:500) for 2 hours at room temperature. Cells were washed again three times using blocking solution, and the secondary antibody (Alexa Fluor goat α-mouse 488 (LI-COR 926-68070); 1:1,000 in blocking solution) was added for 45 minutes at room temperature. After washing again three times in blocking solution, conjugated mouse α*Pf*s25 antibody was added (1:200 in blocking solution) for 30 minutes at room temperature. Cells were washed again three times and mounted using DAPI Fluoromount (Southern Biotech). From each sample, >100 gametocytes were analyzed for quantification.

### *PfGEP1* amplicon sequencing

gDNA was isolated from 200 μl blood using the QIAamp® DNA Blood Mini Kit (Qia-gen) according to manufacturer’s protocols. The region containing the described SNPs was amplified using primers G47ExF and G47ExR (Bustamante et al., 2012; Table S2). All positive samples were Sanger sequenced (LGC Genomics, Berlin) using the amplification primers.

## Author contributions

F.H., A.G.M., and K.M. designed and conceptualized the experiments; F.H., M.S.C., and L.L. performed the experiments; F.K. generated the blood sample collection; F.H., F.K., A.G.M., and K.M. analyzed and interpreted the data; F.H. and K.M. wrote the manuscript; all authors contributed to the article and approved the submitted version.

## Acknowledgments

We thank Pietro Alano (Istituto Superiore di Sanitá, Rome) and Pablo Cortes (Max-Planck Institute for Infection Biology, Berlin) for the kind donation of the anti-*Pf*s25 antibodies.

## Supplemental Information

**Table S1:**
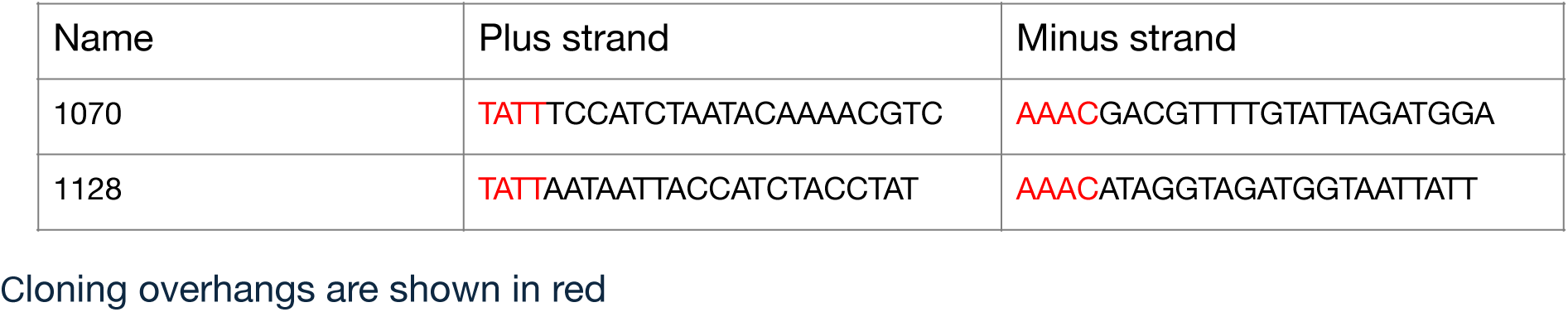
gRNAs.

**Table S2:**
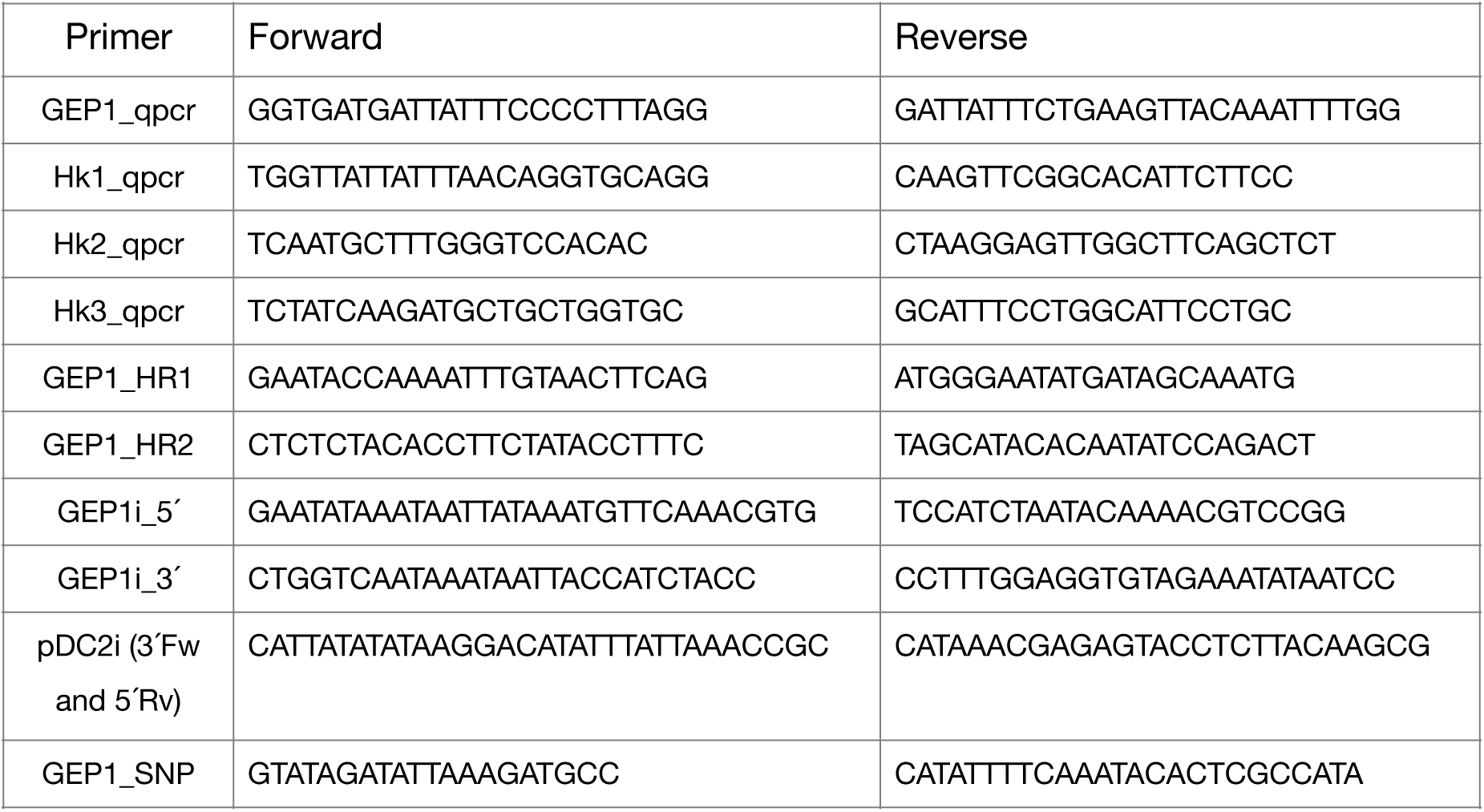
Primers used in this study.

